# The rise and fall of SARS-CoV-2 variants and the mutational profile of Omicron

**DOI:** 10.1101/2021.12.16.473096

**Authors:** Tanner Wiegand, Aidan McVey, Anna Nemudraia, Artem Nemudryi, Blake Wiedenheft

**Affiliations:** Department of Microbiology and Cell Biology, Montana State University, Bozeman, MT 59717, USA

## Abstract

In late December of 2019, high throughput sequencing technologies enabled rapid identification of SARS-CoV-2 as the etiological agent of COVID-19, and global sequencing efforts are now a critical tool for monitoring the ongoing spread and evolution of this virus. Here, we analyze a subset (n=87,032) of all publicly available SARS-CoV-2 genomes (n=∼5.6 million) that were randomly selected, but equally distributed over the course of the pandemic. We plot the appearance of new variants of concern (VOCs) over time and show that the mutation rates in Omicron viruses are significantly greater than those in previously identified SARS-CoV-2 variants. Mutations in Omicron are primarily restricted to the spike protein, while 25 other viral proteins— including those involved in SARS-CoV-2 replication—are highly conserved. Collectively, this suggests that the genetic distinction of Omicron primarily arose from selective pressures on the spike, and that the fidelity of replication of this variant has not been altered.

**Importance:** Omicron is the fifth SARS-CoV-2 variant to be designated a Variant of Concern (VOC) by the World Health Organization (WHO). Here we provide a retrospective analysis of SARS-CoV-2 variants and explain how the Omicron variant is distinct. Our work shows that the spike protein is a ‘hotspot’ for viral evolution in all variants, suggesting that existing vaccines and diagnostics that target this protein may become less effective against Omicron and that our therapeutic and public health strategies will have to evolve along with the virus.

## Main Text

All viruses, including SARS-CoV-2, accumulate mutations as they replicate and spread. Most of these changes have little or no impact on transmissibility, disease severity, or the effectiveness of current vaccines and diagnostics. However, selective pressures that act on random mutations efficiently enrich rare variants with enhanced viral fitness (e.g., replication and transmissibility). Initially, SARS-CoV-2 variants were named according to their geographic origin (i.e., Wuhan), but naming based on geography can be culturally insensitive, stigmatizing, and is often inaccurate. Thus, this naming scheme was quickly replaced by a unique combination of letters and numbers known as the Pangolin nomenclature (e.g., B.1.1.7), but this taxonomic scheme can be cumbersome and sometimes confusing for scientists and the public alike (Rambaut *et al*, 2020). With increasing frequency, we now complement the Pangolin nomenclature with the use of a single Greek letter. Omicron (B.1.1.529) is the 15^th^ letter of the Greek alphabet, but only the fifth variant designated as a Variant of Concern (VOC) by the World Health Organization (WHO).

Omicron was first identified from a specimen collected on November 9^th^ of 2021 in South Africa, and was designated as a VOC on November 26^th^ (WHO, 2021). This designation is based on the number of mutations (26-32) in the spike protein relative to previously sequenced isolates, as well as concerning epidemiological reports from South Africa (Callaway & Ledford, 2021; Chen *et al*, 2021). However, the spike is only one of at least 26 SARS-CoV-2 encoded proteins (Finkel *et al*, 2021). Here we take a step back and provide a general overview of how SARS-CoV-2 genomes have changed over the course of the pandemic. We limit this analysis to the SARS-CoV-2 proteome (i.e., non-synonymous mutations) and highlight how different proteins evolve at distinct rates.

First, we analyze a subset (n=87,032) of all SARS-CoV-2 genomes (n=∼5.6 million) that were randomly selected, but equally distributed over the course of the pandemic; starting with the Wuhan reference genome and ending with all high-quality Omicron genomes that are currently available on GISAID (n=275, on December 4, 2021) (Elbe & Buckland-Merrett, 2017). Overall, this analysis reveals a trend of increasing non-synonymous mutations over time and a significant jump in Omicron, relative to all the previous strains of the virus (**Figure 1A and B**).

**Figure 1.**
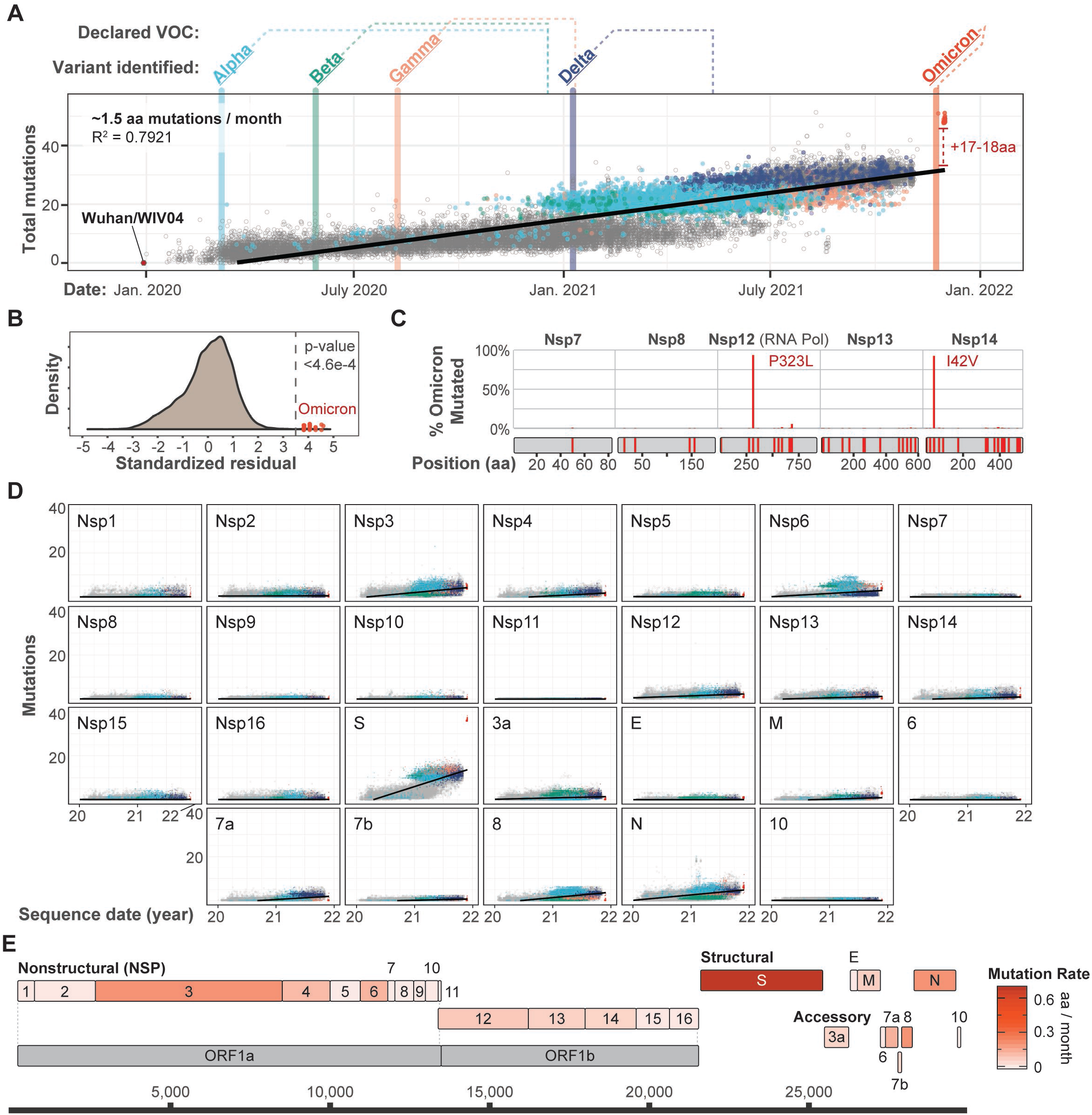
Evolution SARS-CoV-2 and the emergence of new variants. **A)** Mutations acquired over time in 26 SARS-CoV-2 protein sequences extracted from 87,032 genomes (86,757 genomes from Dec 19, 2021 to Nov 10, 2021 and 275 Omicron genomes from Nov 23, 2021 to Dec 1, 2021) (GISAID accessions available at: github.com/WiedenheftLab/Omicron). Variants of concern (VOC) are shown in bold, with dates of the first sequence for each lineage shown as vertical lines through the graph. The time elapsed between first detection and VOC designation by the WHO is shown as dotted lines above the graph. Dots are colored similar to variant names, and grey circles represent non-VOC lineage genomes. A linear regression line is shown in black. Omicron variants deviate from the trend. **B)** Distribution of standardized residuals was plotted to determine how observed mutation numbers in SARS-CoV-2 genomes deviate from the regression line (predicted number) shown in panel A. Omicron genomes deviate >3.5 sigma from the mean, indicating significantly more non-synonymous mutations than expected by the linear model (p < 4.6e-4). **C)** Non-synonymous mutations in the Omicron RNA replication proteins are shown as red lines on schematic representations of each protein, and frequencies of each mutation shown as vertical lines (red). Most non-synonymous mutations are found in less than 1% of Omicron sequences. **D)** Mutations accumulated in SARS-CoV-2 proteins over time. Linear regression lines and variant colors are plotted as in panel A. **E)** Schematic depiction of SARS-CoV-2 protein coding sequences, with each gene colored according to amino acid mutation rates predicted in panel.

Based on the anomalous mutational profiles of Omicron viruses, we hypothesized that Omicron genomes would have mutations that effect the fidelity of viral RNA replication machinery. To test this hypothesis, we analyzed the sequences of replication-associated proteins (i.e., Nsp7, Nsp8, Nsp12, Nsp13 and Nsp14) to determine if they had acquired mutations that would be likely to affect the fidelity of replication. This analysis revealed only one widespread Omicron mutation in the RNA-dependent RNA polymerase (i.e., P323L in Nsp12) (**Figure 1C**). This mutation is expected to have arisen prior to the emergence of the Omicron lineage, since P323L was previously shown to cooccur with a spike mutation (D614G) that swept to global fixation in mid-2020 and is present in 96.3% of all the SARS-CoV-2 sequences we analyzed (Kannan *et al*, 2020; Korber *et al*, 2020). The only other widespread mutation we detected in Omicron RNA replication components was a conservative amino acid substitution (i.e., I42V) in the 3’-5’ exoribonuclease protein (Nsp14) that is thought to be responsible for proof-reading (**Figure 1C**) (Hsu *et al*, 2021). The I42V substitution is distant from the Nsp14 active site (Ma *et al*, 2015), and the P323L mutation in Nsp12 is not unique to Omicron variants. Thus, we do not expect that either of these mutations explain the significant increase in non-synonymous mutations acquired by Omicron lineage viruses.

While the origins and conditions leading to the evolution of Omicron remain uncertain, some scientists suspect that Omicron arose in chronically infected immunocompromised patients where the immune response is too weak to clear the virus but is strong enough to select for mutations that increase viral fitness (Kupferschmidt, 2021). To look for other proteins that might be evolving under similar selective pressures, we evaluated mutational trends for each SARS-CoV-2 protein over the course of the pandemic (**Figure 1D**). This analysis highlights the unique evolutionary signature of the spike protein, which has accumulated non-synonymous mutations more rapidly than any other viral protein—regardless of viral lineage (**Figure 1E**).

Collectively, these data suggest that the spike protein is evolving under particularly strong selection, and this trend is especially pronounced in Omicron variants (**Figure 1D**). More data are necessary to understand how spike mutations impact viral transmissibility, epidemiology, host range and receptor affinity, or clinical severity, but ongoing evolution of the spike is anticipated to erode the efficacy of vaccines designed to target the original Wuhan spike sequence and may complicate the ongoing vaccination campaign (Cele *et al*, 2021; Roessler *et al*, 2021; Wilhelm *et al*, 2021).

